# Virucidal activity of a proprietary blend of plant-based oils (Viruxal) against SARS-CoV-2 and influenza viruses – an *in vitro* study

**DOI:** 10.1101/2021.05.31.446420

**Authors:** Jón Magnús Kristjánsson, Óttar Rolfsson

## Abstract

**Background:** The emergence of a novel coronavirus known as SARS-CoV-2 resulting in a global pandemic COVID-19 has led to a dramatic loss of life worldwide and presented an unprecedented challenge to public health. Viruxal is a medical device in a form of a nasal and oral spay containing a a proprietary blend of plant-based oils which acts against enveloped viruses. The aim of this study was to evaluate the virus deactivation activity of Viruxal against SARS-CoV-2 and Influenza A(H1N1) viruses.

**Methods:** An assay to detect virucidal activity was performed with four concentrations of Viruxal on two virus suspensions. Assessments were made based on log reduction values measured from the assay.

**Results:** Viruxal exhibited virucidal activity by reducing virus titer more than 90% for the enveloped viruses SARS-CoV-2 and influenza A(H1N1) after 30 minutes contact.

**Conclusions:** Viruxal was validated for its potential usefulness as a medical device for treatment and prevention of enveloped respiratory viruses.

## Background

In 2020, the novel corona virus shook the world with its rapid spread and lethal consequences. COVID-19, caused by SARS-CoV-2 virus, became a global pandemic and posed as a huge threat to healthcare systems in many countries^1^. Currently, there is no effective cure for COVID-19. The disease spreads by person-to person transmission and can lead to severe illness and even death. The SARS-CoV-2 virus is transmitted directly through airborne respiratory droplets and indirectly from exposed surfaces. The virus enters the body through the mouth, nose and eyes, attaching to the mucosal tissue where it enters the mucosal cells and multiplies^2^. In recognition of the transmission mechanism of SARS-CoV-2 virus, official guideline and safety measures were rapidly established. Safety measures such as usage of facial masks, frequent handwashing and social distancing are scientifically proven to reduce the spread of COVID-19 in public^3,4^. However, there are some situations where the risk of infection is particularly high, such as public transportation, international travel or in crowded areas or where these safety guidelines are impractical to follow, such as eating in restaurants or participating in certain types of sports activities. In these situations, alternative or additional precautions approaches may be beneficial.

Major common respiratory viruses include influenza, rhinovirus, human coronaviruses, RSV, parainfluenza and adenoviruses^7,8^. Most of these viruses initially infect the upper respiratory tract, thus creating a topical treatment for the nasopharynx would be beneficial. Researchers are further concerned about the synergistic effects of a coinfection with SARS-CoV-2 and other viruses^5,6^, especially influenza.

Natural oils can form a preventative surface barrier for viral infections through an arrangement of positively and negatively charged fatty acids. Viruxal is administered onto the mucosa of the nasal/oral cavity to reduce microbial adhesion and deactivate and inhibit virus filled droplets from infecting mucosal tissues. The device has demonstrated its safety and effectiveness from a series of *in vitro* testing and ongoing clinical trials. In the study presented here, the virucidal effects of Viruxal against the SARS-CoV-2 and Influenza A(H1N1) viruses were investigated by titrating Viruxal onto viral populations and assaying the effect on viral load. The results indicate that Viruxal has potent virucidal effect and thus could potentially be used as additional protection for the nasal and oral cavity against these viruses.

## Methods

### Test substance

The test substances were; ViruxOral and ViruxNasal Sprays, manufactured by Viruxal-Iceland. ViruxOral and Nasal are identically formulated with regards to active ingredients and collectively referred to as Viruxal in this study. The mode of action is based in virucidal activity of natural fatty acids (oleic acid, palmitic acid, linoleic acid, stearic acid, linolenic triglycerides and free fatty acids), the main ingredients of the spray. The spray was tested undiluted, (100%), 50%, 25% and 12.5%.

### Virus, Media and Cells

SARS-CoV-2 virus (USA-WA1/2020) was obtained from a patient with laboratory-confirmed COVID-19 in January 2020 in Washington, USA and inoculated in Vero 76 cells cultured in Minimum Essential Medium (MEM) (HyClone, Cytiva) supplemented with 2 % fetal bovine serum (FBS) (HyClone, Cytiva) and 50 μg/mL gentamicin (Sigma). Human influenza A virus strain, A/California/07/09 (H1N1) pdm09 (International Reagent Resource, IRR) was grown in Madin-Darby canine kidney (MDCK) cells with serum-free MEM (HyClone, Cytiva) containing 1 IU/mL of trypsin (Sigma), 1 μg/mL EDTA (Sigma) and gentamicin (Sigma). To determine the virus titer, the respective cells were incubated with serially diluted virus stock solutions. The selected virus titer were the concentrations that yield an observable amount of cytopathic effects (CPE) during the incubation period. The study was executed by an independent university laboratory in the United States.

### Virucidal Assay

Virucidal assays are used to determine if a compound inactivates virus outside of cells, or in other words, whether the compound deactivates the virus before it infects the host cells. The assay is performed by incubating the virus and testing compounds for a pre-specified time before inactivation, followed by a determination of virus titer using dilution assay in 96-well microplates of cells calculated by the Reed-Muench method^7^.The Reed Muench method measures the virus titer required to either kill 50% of cells (cell culture infected dose, CCID_50_) or to produce a cytopathic effect (CPE) in 50% of inoculated culture cells. At the end of incubation period, the log reduction values (LRV) of testing compounds is calculated and compared against the negative control (water).

Viruxal was tested at four concentrations (100, 50, 25 and 12.5%) against the SARS-CoV-2 or influenza A (H1N1) viruses. Compounds and the respective virus were incubated at room temperature for three contact times of 1, 5, and 30 minutes. Virus stocks were added to triplicate tubes of each prepared concentration so that there was 50% virus solution by volume and 50% prepared sample. Ethanol was tested in parallel as a positive control and water as the negative control. In addition, media alone was added instead of virus to one tube of each prepared concentration to serve as toxicity control. Following the contact period, the solutions were neutralized by a 1:10 dilution in test media and each dilution was added to 4 wells of a 96-well plate with 80-100% confluent cells. The cells were incubated at 37 ± 2°C with 5% CO2. On day 5-7 post infection, the plates were observed for CPE, calculated for CCID_50_ and LRV.

### Controls

Virus controls were tested in water and the reduction of virus in test wells compared to virus controls were calculated as the log reduction value (LRV). Toxicity controls contained Viruxal and media in the absence of virus. Neutralization controls were tested to ensure that virus inactivation did not continue after the specified contact time, and that the residual sample in the titer assay plates did not inhibit growth and detection of surviving virus. This was done by adding toxicity samples to titer test plates then spiking each one with a low amount of virus that would produce an observable amount of CPE during the incubation period.

## Results

To determine the virus deactivating capability of the Viruxal spray on different virus strains, Viruxal was incubated at four concentrations with the SARS-CoV-2 or influenza A(H1N1) virus stocks as described in the methods. Samples from each incubation were titrated with the 50% Cell Culture Infectious Dose (CCID_50_) using the appropriate host cell systems for each virus.

### 1. Viruxal inactivates SARS-CoV-2 by 95 % independent of treatment time

Virus titer and log reduction values of the Viruxal compound tested against SARS-CoV-2 were recorded and shown in **Table 1**. Viruxal exhibited virus inactivation activity by reducing the virus titer more than 1 log for SARS-CoV-2. After a 1-minute contact time, Viruxal reduced SARS-CoV-2 by >1 log, equivalent to >90% inactivation of the virus, in concentrations from 25-100%. After 30 minutes, Viruxal lowered SARS-CoV-2 virus by >1 log (>90%) at all tested concentrations and by >2 logs, equivalent to >99% inactivation of the virus, at 50, 25 and 12.5% concentrations. SARS-CoV-2 virus was not reduced below the limit of detection at any concentration regardless of contact time (**Figure 1**).

**Table 1.**
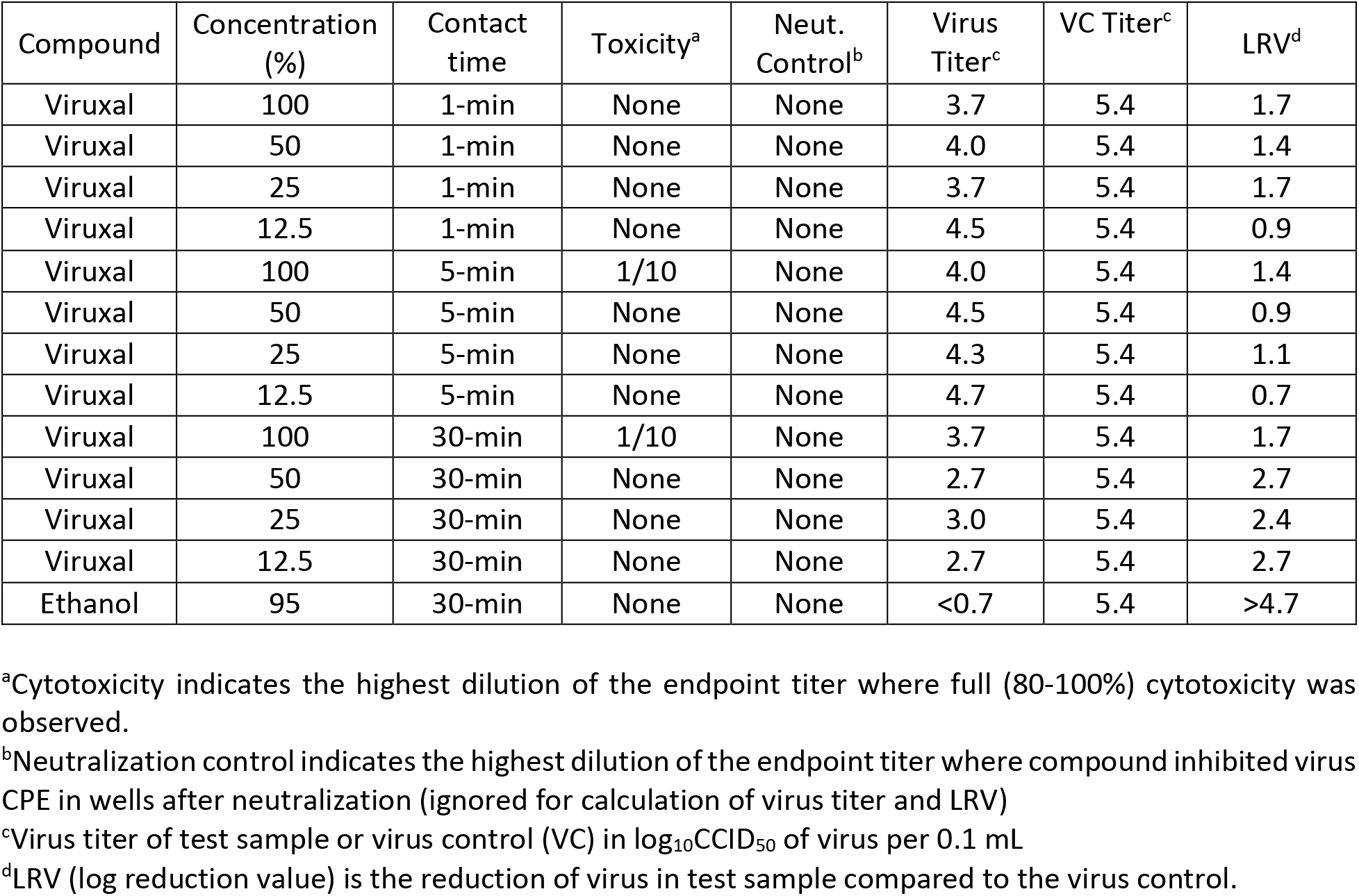
Virucidal efficacy against SARS-CoV-2 after incubation with virus at 22 ± 2°C.

**Figure 1.**
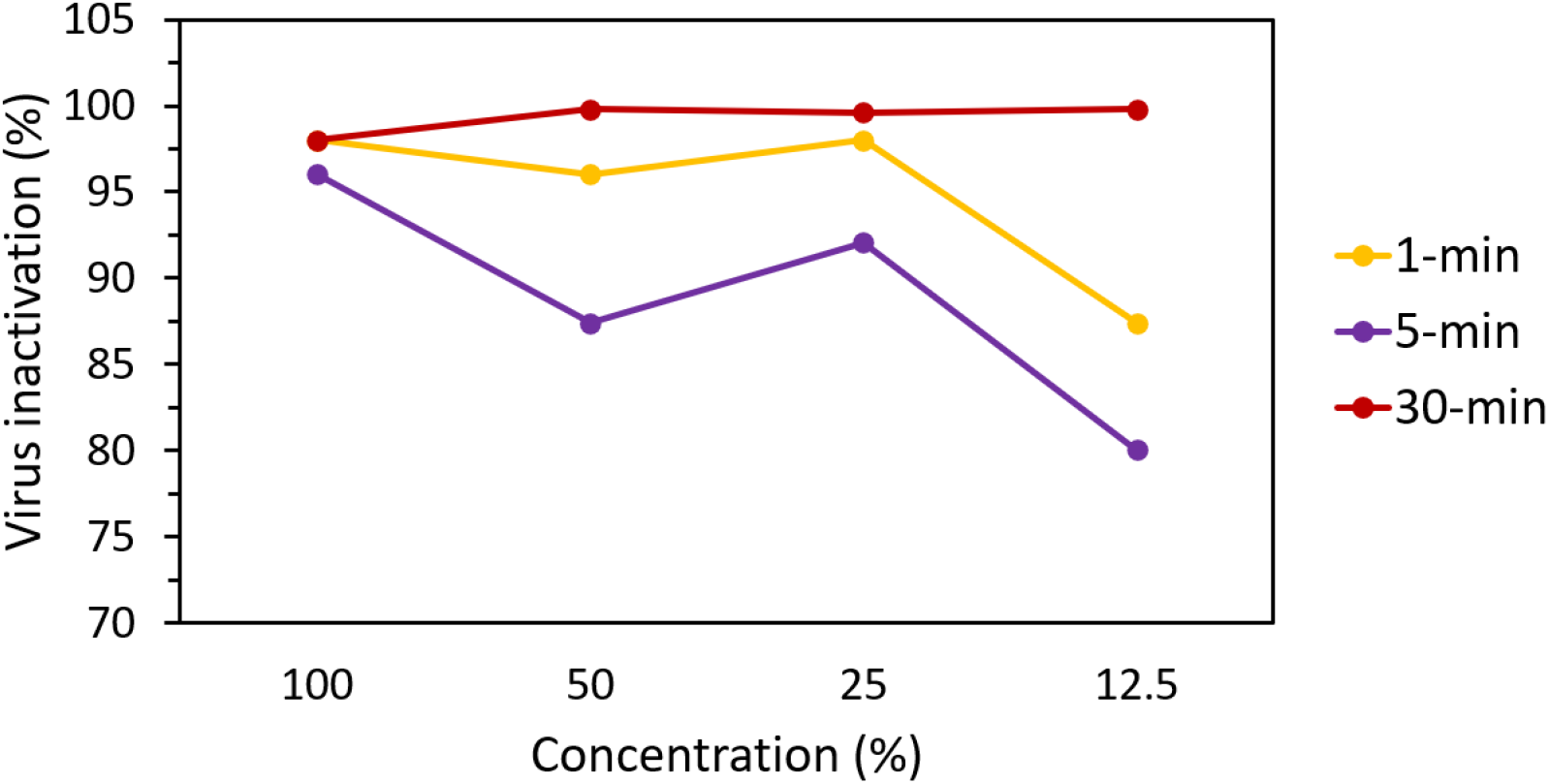
SARS-CoV-2 virus inactivation after 1 and 30 minutes at different concentrations.

### 2. Viruxal inactivates Influenza A(H1N1) by 99 % independent of treatment time

Influenza A(H1N1) virus titers were reduced by >1 log (>90%) at all tested concentrations and time points (Table 2). At 1-minute contact, 100% Viruxal yielded more than 3 logs reduction corresponding to 99.9% virus inactivation and 50% Viruxal deactivated more than 99% (>2log) viruses. After 5-minute contact, the full and half concentrated Viruxal resulted in a more than 2 logs reduction (>99%). At 30-minute incubation, all the concentrations showed a greater than 2 logs virucidal effects, and a greater than 3 logs at 100% concentration. Virus was not reduced below the limit of detection (**Figure 2**).

**Table 2.** Virucidal efficacy against influenza A(H1N1) after incubation with virus at 22 ± 2°C.

| Compound | Concentration (%) | Contact time | Toxicity<sup>a</sup> | Neut. Control<sup>b</sup> | Virus Titer<sup>c</sup> | VC Titer<sup>c</sup> | LRV<sup>d</sup> |
| --- | --- | --- | --- | --- | --- | --- | --- |
| Viruxal | 100 | 1-min | None | None | 3.0 | 6.2 | 3.2 |
| Viruxal | 50 | 1-min | None | None | 3.5 | 6.2 | 2.7 |
| Viruxal | 25 | 1-min | None | None | 4.7 | 6.2 | 1.5 |
| Viruxal | 12.5 | 1-min | None | None | 4.3 | 6.2 | 1.9 |
| Viruxal | 100 | 5-min | None | None | 3.7 | 6.2 | 2.5 |
| Viruxal | 50 | 5-min | None | None | 4.0 | 6.2 | 2.2 |
| Viruxal | 25 | 5-min | None | None | 4.3 | 6.2 | 1.9 |
| Viruxal | 12.5 | 5-min | None | None | 4.5 | 6.2 | 1.7 |
| Viruxal | 100 | 30-min | None | None | 2.4 | 6.2 | 3.9 |
| Viruxal | 50 | 30-min | None | None | 3.5 | 6.2 | 2.7 |
| Viruxal | 25 | 30-min | None | None | 3.7 | 6.2 | 2.5 |
| Viruxal | 12.5 | 30-min | None | None | 3.5 | 6.2 | 2.7 |
| Ethanol | 95 | 30-min | None | None | <0.7 | 6.2 | >5.5 |

<sup>a</sup>Cytotoxicity indicates the highest dilution of the endpoint titer where full (80-100%) cytotoxicity was observed.
<sup>b</sup>Neutralization control indicates the highest dilution of the endpoint titer where compound inhibited virus CPE in wells after neutralization (ignored for calculation of virus titer and LRV)
<sup>c</sup>Virus titer of test sample or virus control (VC) in log<sub>10</sub>CCID<sub>50</sub> of virus per 0.1 mL
<sup>d</sup>LRV (log reduction value) is the reduction of virus in test sample compared to the virus control.

**Figure 2.**
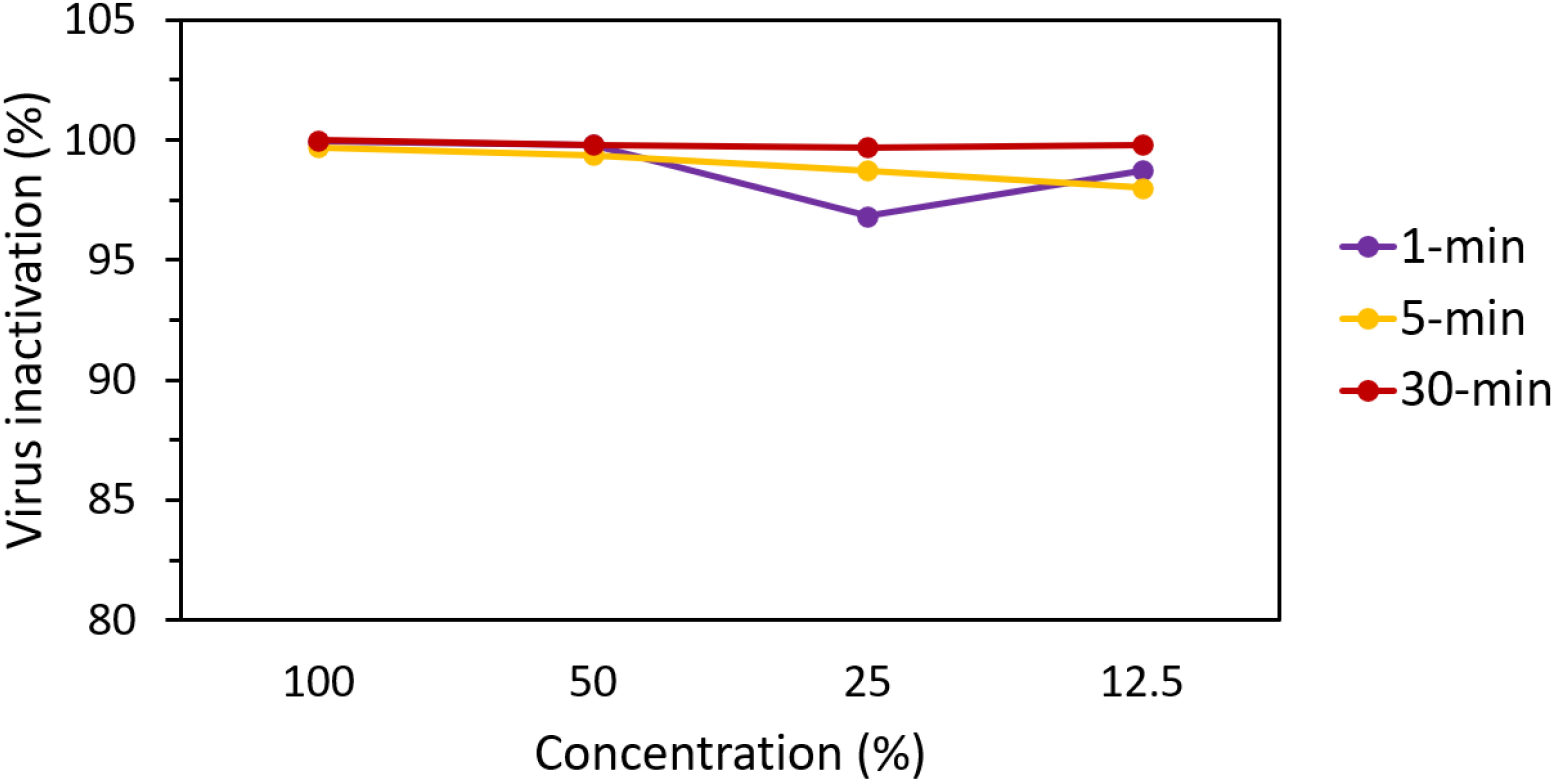
Influenza A(H1N1) virus inactivation after 1 and 30 minutes at different concentrations.

Neutralization controls demonstrated that residual sample inhibited virus growth and detection of SARS-CoV-2 and influenza viruses in the endpoint titer assays in some wells, though never in both neutralization test wells and so it was not noted in **Tables 1–2** and the effect was ignored for calculation of titers and LRV. Positive controls performed as expected.

## Discussion

The present study demonstrates strong *in vitro* viral inactivation efficacy of the fatty acid based product against the enveloped viruses SARS-CoV-2 and influenza A(H1N1). A variety of Viruxal concentrations, ranging from 12.5 to 100%, were tested and showed virucidal effect with 80% to 99.9% viral deactivation at an incubation time of 1-30 minutes. The results suggest that Viruxal may be active against major enveloped viruses causing respiratory diseases, including the one causing the global pandemic of COVID-19.

The basic mechanism for the virucidal activity of the Viruxal technology is to deliver a potent fatty acid layer to the oral and nasal cavity the primary site of respiratory virus infection and replication. When applied topically on the oral and nasal mucosal tissue, it is hypothesized that the fatty acid molecules in Viruxal, penetrate and break down the lipid coating that covers enveloped viruses, thereby deactivating the virus at their point of entry. The antibacterial and antiviral effects of fatty acids have been studied and reported extensively in the literature. Fatty acids found in nature in animal and vegetable fats, milks and oils are known to possess inhibitory activities towards bacteria, fungi, yeast and enveloped viruses. Linoleic acid, linolenic acid and to a lesser degree oleic acid exhibit bacteriostatic effects on gram-positive and gram-negative bacteria^9^. Both unsaturated fatty acids oleic and linoleic have been shown to deactivate enveloped viruses such as herpes, influenza, Sendai and Sindbis within a short contact time^10^. Fresh human milk contains a rich amount of fatty acids and has been shown to protect infants from viral infections. In fact, Thormar *et al* demonstrated that the fatty acids in milk lipids expressed antiviral activities against several enveloped viruses, especially when the milk was stored at 4°C^11^. Further experimental evidence has demonstrated that medium-chain saturated and long-chain unsaturated fatty acids as present in Viruxal are highly active while short-chain and long-chain saturated fatty acids had no or very little antiviral activities^11–14^. The antiviral mechanism of fatty acids has been elucidated through the disruption of viral envelopes, causing leakage and potentially a complete disintegration of the envelope and the viral particles (figure 3)^11^. Even though the exact mechanism in which Viruxal fatty acids disrupts the virus membrane requires further investigation, it can be hypothesized that the number of chains and saturation level of the fatty acids in the spray directly affect the viral membrane stability and therefore its infiltration proficiency.

**Figure 3.**
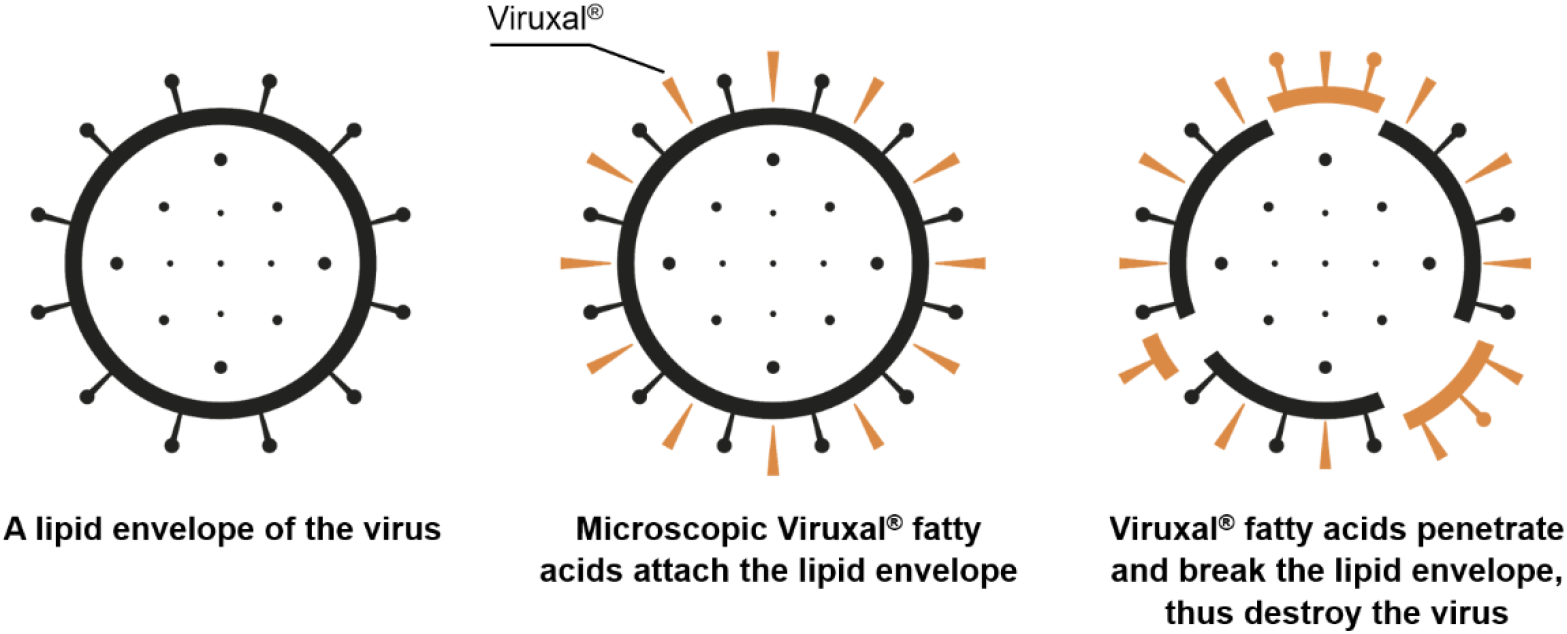
A proposed antiviral mechanism of fatty acids in Viruxal against enveloped viruses.

Since the first SARS-CoV-2 virus was identified in Wuhan, China in December 2019. The 2019 novel coronavirus has swept across multiple continents and become a Public Health Emergency. As of January 22, 2021, approximately 10 months after the COVID-19 outbreak was declared a pandemic, there have been more than 96 million confirmed cases and over 2 million deaths recorded globally according to the World Health Organization. While two vaccines have already received an Emergency Use Authorization (EUA) by the US Food and Drug Administration (FDA), the vaccine distribution plan will take months, perhaps even years to reach populations in developing countries. Furthermore, seasonal influenza and other respiratory viruses can intensify and worsen the outcomes of COVID-19, hindering the pandemic control. In the search for treatments for COVID-19, many researchers focus on the mechanism of SARS-CoV-2 virus infecting human cells. SARS-CoV-2 is an enveloped, single stranded RNA virus that enters human body via binding to angiotensin-converting enzyme 2 (ACE2). ACE2 is a protein on the surface of many cell types and especially abundant in epithelium in the nose, mouth and lungs^15^. When SARS-CoV-2 virus binds to ACE2, it suppresses the protein’s functions and causes damage in tissues with high expression of ACE2, especially the lung alveoli, leading to acute lung damage, respiratory distress syndrome and pneumonia^16^. Higher virus shedding links to more severe COVID-19 symptoms and complications^17^. Sine the nasal and oral cavity is home to virus seeding, replication and ACE2 localization, lowering the local viral load can potentially lessen symptoms and reduce the risk of transmission. Viruxal delivers a fatty acid layer to cover the prone nasal and oral cavity, which deactivates the virus and diminishes the amount of virus entering the body by preventing it from binding to cellular receptors. Also, the application of Viruxal on infected subjects could decrease the viral load in expelled virusols, hence lowering the risk of spreading virus to the community.

The virucidal effect of Viruxal against Influenza A(H1N1) in 30-minute incubation was very encouraging. Viruxal yields more than 99% virucidal efficacy at all four tested concentrations. Similar to SARS-CoV-2, Influenza A(H1N1) primarily targets the human respiratory system and results in mild to severe illness, acute pneumonia and even respiratory failure. Seasonal influenza occurs annually, especially in the winter, with the fatality rate of 0.1-0.2%. Scientists have noticed coinfections between influenza A and SARS-CoV-2 to cause more severe disease and an increased risk of death than infection by either virus *in vitro* and clinically^5,18,19^. Therefore, applications of Viruxal not only protect patients from the two viruses but also could potentially reduce the risk of dangerous coinfections. Our study was performed *in vitro* and limited to single virus infections; further studies are needed to understand the mechanism of coinfection suppression capability of Viruxal.

## Conclusions

In summary, the results show potent viral deactivation of the proprietary blend of plant-based oils in the Viruxal medical device against SARS-CoV-2 and Influenza A(H1N1). Although additional research and clinical evidence are needed to further support the widespread use of Viruxal in a clinical setting, this study implies the potential of Viruxal as a prevention approach and a possible therapeutic treatment and for major respiratory infections caused by encapsulated viruses.

## Ethics approval and consent to participate

Not applicable

## Author’s contributions

All authors have contributed to acquisition, analysis or interpretation of data; writing or editing of the manuscript. All others read and approved the final manuscript.

## Availability of data and materials

The datasets analyzed during the current study are not publicly available but are available from the corresponding author on reasonable request.

## Acknowledgement

Not applicable

## Consent for publication

Not applicable

## Competing interests

The authors declare that they have no competing interests.

**Authors’ information**

